# A Long Timescale Stimulus History Effect in the Primary Visual Cortex

**DOI:** 10.1101/585539

**Authors:** Hyewon Kim, Jan Homann, David W. Tank, Michael J. Berry

## Abstract

Responses of neurons in the primary visual cortex (V1) are often understood as encoding the current visual stimulus. Yet, some studies indicate that temporal contingency effects exist in the responses of neurons in early sensory areas. We explored if the recent stimulus history would alter the response of V1 layer 2/3 pyramidal cells in head-fixed awake mice during presentation of sequences of complex images. The activity of individual neurons was sparse, such that either one or none of the images in the sequence typically yielded a strong response. We then substituted an image preceding this primary image in order to determine if responses to the primary image were affected. We found that the amplitude of the neuron’s response could be significantly altered by substitutions up to five images back from the primary image, even when the substituted image elicited virtually no response by itself. This stimulus history effect was heterogeneous across the population, with some cells showing facilitation and others suppression. For individual cells, the history effect was robust and reproducible across days. Our data show that responses of V1 neurons not only reflect the current stimulus but also encode, through their response amplitude, information about multiple images previously presented as far as 1000 msec in the past. This might enable V1 to retain information about the extended trajectory of past stimuli and perform complex temporal computations that are as of yet not appreciated.

## 1 Introduction

A constantly changing stream of information enters the visual system at any moment in time. Yet, many experiments have focused only on the responses of visual neurons to temporally isolated stimuli. Some examples exist, however, of altered responses when the temporal context of the stimulus is changed. For example, repeated presentation of an image in macaque inferotemporal cortex (IT) causes a smaller response to the second presentation [1], a phenomenon called repetition suppression. Furthermore, learned transitions from one image to another also cause a strong suppression of the response to the second image, a phenomenon called prediction suppression [2]. In addition, deviant stimuli in a series of common stimuli have been shown to elicit larger responses in IT compared to the common stimulus [2].

In addition to history effects high in the sensory hierarchy, it is increasingly becoming appreciated that early sensory areas may also exhibit forms of memory that exceed commonly understood integration times. In the primary auditory cortex (A1) of the cat, there is evidence for multiple timescales of processing leading to the extraction of auditory objects [3][4][5][6][7]. In A1 of ferret, responses contain information about the previous tone [8]. In the barrel cortex (S1) of the mouse, responses of cells in the supragranular layers contain information about the time interval between whisker stimuli [9]. Stimulus history effects have even been found in the somatosensory thalamus, where the adaptation of a response contains information about a previous whisker stimulus [10].

Similar to repetition suppression in IT, contrast adaptation in V1 decreases the response of neurons to the second presentation of the same stimulus [11][12]. Furthermore, this phenomenon can be specific to the orientation, spatial frequency, and other properties of the stimulus [13][14][15]. V1 neurons in mice have been reported to show an increased response to a deviant stimulus, above and beyond the effects of stimulus-specific adaptation [16]; however, another study in rat using a more diverse set of images did not find an explicit deviance response [17]. Extensive studies have been conducted in A1 to tease apart the temporal contingency phenomena of mismatch negativity, which exhibits true deviance detection, from stimulus-specific adaptation [4][18][19]. In any case, deviance detection – the neural substrate for which has not entirely been elucidated as of yet – is known to be dependent on stimulus history [20], as is stimulus-specific adaptation. Furthermore, it has been shown in the marmoset monkey that a second tone is significantly modulated by the presence of the first, which suggests that auditory responses in the cortex are sensitive to stimulus context [21].

Most of these studies have found memory effects on quite short time scales. However, longer time scales were found in a study in the primary visual cortex of the anesthetized cat [22]. Here, individual static stimuli or short sequences of stimuli were presented and neural activity of 100+ neurons was used to decode stimulus identity at different points in time. The authors found above-chance performance extending up to 700 msec after pre-sentation of individual stimuli and 400 msec after stimuli within short sequences. This result indicated the presence of a form of short-term memory at the population level, and was interpreted as evidence in favor of liquid-state dynamics [22][23]. Questions yet outstanding from this study include how individual neurons encode stimulus history and whether similar effects might be seen in awake animals.

With this background in mind, we wanted to explore temporal contingency effects early in the visual hierarchy over long time scales and for awake animals. We recorded calcium signals simultaneously from 50-100 neurons in the primary visual cortex of awake mice and presented sequences of complex images. In order to assess whether neural activity reflected stimulus history, we substituted images in the sequence and measured the change in response to subsequent images. We found that the responses of layer 2/3 excitatory neurons were significantly modulated by the history of preceding stimuli, even when such preceding stimuli did not elicit a response by themselves. We found effects reaching up to 5 images back in the sequence, which corresponded to a history effect extending up to 1000 msec into the past. These results suggest that a general short-term memory mechanism exists in the primary visual cortex in awake mice.

## 2 Methods

### 2.1 Animal surgery and husbandry

We performed all experiments according to the Guide for the Care and Use of Laboratory Animals, and all procedures were approved by Princeton University’s Animal Care and Use Committee. We used male C57BL/6 mice from the transgenic mouse line Emx1-Cre Ai93 (*n* = 4 mice), which have widespread expression of the calcium indicator GCaMP6f in excitatory neurons of the neocortex, striatum, and hippocampus [24][25]. Recordings were performed on the mice when they were 4-8 months of age.

For the purpose of imaging, we implanted a chronic cranial window above the visual areas of each mouse. We followed the same surgical procedure as outlined in Homann *et al*. [26], adapted from Dombeck *et al*. [27]. Prior to surgery, we administered Dexamethasone at 2 mg/kg to prevent brain swelling. Anesthesia was induced in the mouse with 2.5% isoflourane and maintained with 1.5% isoflurane. We then drilled a 3 mm round craniotomy centered at 2.7 mm lateral and 1.3 mm anterior of lambda with a dental drill. The extracted piece of bone was then substituted with a 3 mm round microscopy cover glass, glued to a thin metal ring and fixed in place with n-butyl cyanoacrylate tissue adhesive (Vetbond, 3M). During this procedure, the dura was left intact. We then attached a titanium headplate to the skull with dental cement (Metabond, Parkell), which we used to fixate the mouse head under the microscope. In addition, a flat, round titanium light shield was attached using Metabond for imaging purposes. Following surgery, we administered the pain reliever Meloxicam at 1 mg/kg and allowed mice to recover for several days before imaging. The mice were then placed on a reversed day-night cycle.

### 2.2 Visual area mapping

We performed widefield one-photon calcium imaging to determine the boundaries of V1. For this, we head-fixed the mouse on top of an air-suspended Styrofoam ball, and surrounded the mouse with a Styrofoam toroidal projection screen. To prevent light leakage, we used a custom-made shield constructed out of a black rubber balloon to cover the space between the objective and headplate. Following Zhuang *et al*. [28], we presented a 20-degree wide bar containing a flickering, full-contrast checkerboard pattern. This bar was moved slowly (9 degrees/s) in each of the four cardinal directions [28] over 20 repeated trials. During this presentation, the GCaMP6f fluorescence activity within the full cranial window was imaged at 20 Hz through a tandem-lens low magnification one-photon setup (1.0x dry objective). A blue (470 nm) LED (Luxeon star) provided the one-photon excitation. The power used to drive the LED was 2.5 mW/cm^2^ at the focal plane. The moving bar generated a faint moving wave of calcium activity in every visual area according to their retinotopy, including the primary visual cortex.

For processing, the brain motion caused by the running of the mouse was corrected. To compute the time-averaged fluorescence image, the custom algorithm identified and excluded the pixels falling under a threshold determined by the median intensity of their neighbors. These sub-threshold pixels were determined to be vessels. The ∆*F/F* was then calculated to create retinotopic maps. These azimuthal and altitudinal retinotopic maps informed the best location within V1 to record from later during two-photon calcium imaging. This position in the retinotopic map corresponded to where the stimulus fell within the visual field sensitive to the retina of the mouse. Boundaries between visual areas, identified by reversals in the retinotopic map, were determined as follows with MATLAB code kindly provided by the group of Edward Callaway [29]. First, both retinotopic maps were smoothed using a 2D Gaussian filter function. Then, the spatial gradient of each map was calculated using a noise-robust estimation method [30]. A single visual field sign (VFS) map was constructed by calculating the sine of the angle between the gradients in the azimuthal versus altitudinal maps at each spatial location. Individual visual areas were then determined by contiguous pixels of the same sign. Some adjacent visual areas had the same VFS (for instance, areas V1 and AM). This situation could be identified by calculating the total visual field coverage of a putative cortical area and finding that the coverage exceeded a threshold of 1.1. In this case, the putative area was subdivided into two areas by finding the boundary where the spatial gradients in both the azimuthal and altitudinal maps jumped discontinuously in their direction. Finally, local blood vessels were used as landmarks in order to navigate to the desired field of view in our high magnification two-photon microscopy setup (Fig. 1A).

**Figure 1:**
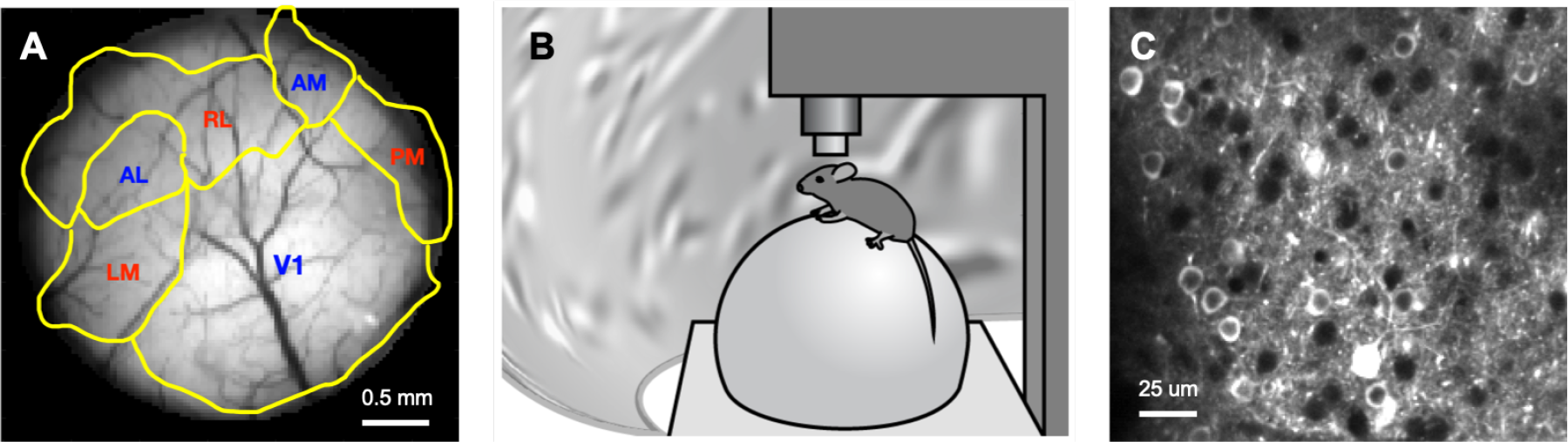
Experimental overview. (A) One-photon image of 3 mm cranial window with visual areas delineated in yellow. Blood vessels are visible (dark); scale bar: 0.5 mm. (B) Two-photon calcium imaging setup with head-fixed mouse on air-suspended Styrofoam ball surrounded by a toroidal projection screen. (C) Partial field of view under our two-photon setup; scale bar: 25 *µ*m.

#### 2.2.1 Two-photon calcium imaging

For our main recordings, we again head-fixed the mouse on an air-suspended Styrofoam ball, and surrounded the mouse with a custom Styrofoam toroidal projection screen (Fig. 1B).

Visual stimuli were projected on this toroidal screen, as previously described [31]. One advantage of the toroidal screen is that it covers a large portion of the field of view of the mouse (−20 to +70 degrees vertically and −130 to +130 degrees horizontally). Such a wide coverage ensures that the position of most neurons’ receptive fields is within the range of the screen.

We then used a two-photon microscope (Titanium sapphire laser, 140 fs pulses at 80 MHz, 920 nm wavelength, water immersion objective, 40x, NA 0.8) to image activity of neurons at cellular resolution at a depth of 150 *µ*m (layer 2/3) in a 410 × 410 *µ*m field of view. This allowed us to record from roughly 100 cells in parallel per session. We used a laser power of 55 mW in all our recording sessions. Fluorescent signals were detected by a photomultiplier tube and processed digitally. Images of 512 × 512 pixels were created by a resonant galvanometer scan mirror system controlled by ScanImage 5.0 (Vidrio Technologies) and acquired at a 30 Hz frame rate. Voltages from the vertical scan mirror were routed to and digitized with a Digidata 1440A (Axon instruments) and recorded in Clampex (Axon instruments). Along with stimulus event markers, these signals were used to align the two photon imaging data with the visual stimulus.

### 2.3 Data processing

We constructed ∆*F/F* traces for individual neurons according to the following procedure:

1. *Images were motion corrected*. Because our animals were awake and were able to move freely on the air-suspended Styrofoam ball, the brain showed slight lateral movements. We corrected those lateral movements with custom scripts in Matlab and ImageJ. This was done by first generating a low noise template by averaging 1000 images during a recording session, and then registering all of the images in that recording session to this template by maximizing the cross-correlation. This process was iteratively repeated 3 times.
2. *Region of interest (ROIs) were selected* (Fig. 1C). Cell somas were traced semi manually based on time averaged frames with the help of the Cell Magic Wand Tool (Max Planck Florida) for ImageJ. With this method we selected neurons independent of their activity, therefore avoiding an activity bias. Somas were identified by their characteristic ‘donut’ shape, which is due to the nuclear exclusion of the calcium sensor. A Matlab code then averaged all pixel values in an ROI, constructing a fluorescence trace across the series of images for each cell.
3. ∆*F/F was computed*. We estimated the baseline of the fluorescence trace *F*_0_(*t*) using the method of [27]. In brief, we divided the fluorescence activity across the entire recording into 15-sec segments. Then, we computed the 8th percentile of the distribution of the fluorescence signals in each segment, which represented our estimate of the average baseline activity in that segment. Finally, we formed a smooth estimate of the baseline *F*_0_(*t*) at the temporal resolution of our raw data by using spline interpolation. This dynamic estimation of the baseline *F*_0_(*t*) ensures that small, slow drifts in fluorescence are accounted for when computing the relative change in fluorescence, ∆*F/F* = [*F* (*t*) – *F*_0_(*t*)] / *F*_0_(*t*).
4. *Stimulus event markers were synched to image acquisition*. Event markers (sampled at 2000 Hz) were then recomputed to match the sampling rate of the image acquisition (sampled at 30 Hz).

#### 2.3.1 Stimulus construction

Each image was composed of a superposition of 100 Gabor patches of random phase, location, orientation, spatial frequency, and spatial extent (Fig. 2) [26]. Parameters of the Gabors were designed to be in the range of measured receptive field parameters of V1 neurons in the mouse [32], ensuring maximal responsiveness of the neurons. Each image in our ensemble had a similar global luminance (+/- 1.2 % across the image ensemble) and contrast, a design that minimized changes in overall activity due to differences in global luminance or contrast.

**Figure 2:**
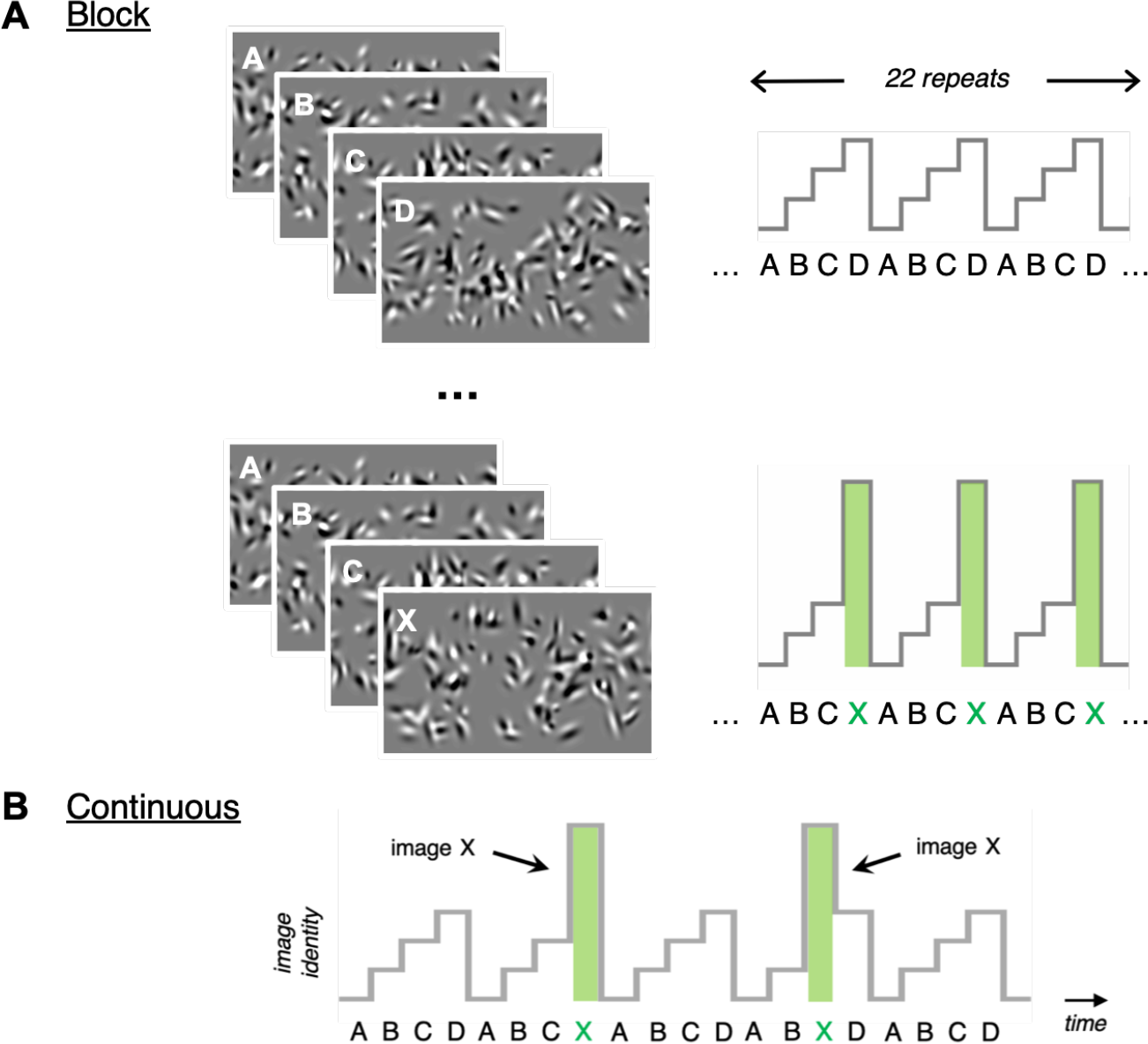
Stimulus design. (A). In the “block” stimulus design, a given sequence was repeated 22 times. There were 5 different image sequences: *ABCD*, *XBCD*, *AXBC*, *ABXD* or *ABCX*. Each image contained a random superposition of 100 Gabor patches. (B) In the “continuous” stimulus paradigm, a sequence of images *A*, *B*, *C* and *D*, was repeated many times and any given image was occasionally (average of once every 5 seconds) substituted with a 5th image *X* drawn from the same image ensemble.

We then selected 4 randomly chosen images from the ensemble described above, denoted *A*, *B*, *C*, *D*, and presented them in a temporal sequence, denoted *ABCD*. In order to test for history effects, we substituted a new image, *X*, for one of the images in the core sequence. The image *X* was randomly drawn from the same ensemble as the other images, so that it was statistically the same but different in detail. With this choice, we altered the detailed stimulus history without causing overall changes in neural activity. We made these substitutions using two different stimulus designs:

1. In the ‘block’ design (Fig. 2A), we repeated a given sequence 22 times in a row. There were five different choices of sequences. There could be either a core sequence, *ABCD*, or one of four substitution sequences. The four substitution sequences were constructed by substituting one of the four images with another randomly chosen image, *X*, denoted by *XBCD*, *AXCD*, *ABXD* or *ABCX*. Each of these five block types were randomly interleaved within a one-hour presentation time, which resulted in 30 presentations of each block type. We also performed two-hour recording sessions, which resulted in 60 presentations of each block type.
2. In the ‘continuous’ design (Fig. 2B), a core sequence, *ABCD*, constructed out of 4 randomly chosen images, was repeated for one or two hours. Individual images were presented for 250 msec each. Within this sequence, images were randomly substituted with a 5th image *X* at a rate of 1 substitution every 5 seconds.

We also carried out experiments using the block design with 8- and 16-image sequences, to test for longer history effects. Within a block, each 8-image sequence repeated 20 times, and 16-image sequence was repeated 10 times. A recording session of one hour then resulted in 17 presentations of each block for both 8- and 16-image sequences. In the case of the 16-image sequence, we restricted substitutions to only odd images, *A*, *C*, *E*, *G*, *I*, *K*, *M* and *O*. This was done so that we could collect enough repeats of each substitution to provide sufficient statistical power.

Occasionally we repeated the same stimulus for 2 or 3 days, recording from the same neurons. This was done in order to test the day-to-day robustness of history effects observed in individual neurons. In all experiments, animals viewed the stimulus passively.

### 2.4 Data analysis

#### 2.4.1 Selection of cells for analysis

We were only interested in cells that responded robustly to at least one image of the core sequence. To this end, we selected cells with a strong sustained response, which was defined as cells with a mean activity of at least 0.2 ∆*F/F* [26]. We also excluded cells that responded to the substitution image *X*, because this response interfered with our analysis of sequence selectivity (see below). *X*-selective cells were defined as those cells that had a strong sustained response that was present in all four substitution sequences but not in the core sequence. In total, these selection criteria resulted in 151 viable neurons out of 1275 recorded cells (11.8%).

#### 2.4.2 Determination of the primary image

We then determined the amplitude of the response of the individual cells to each image. For this, we presented the 4 core images *A*, *B*, *C* and *D* plus the one substitution image *X* in random order for 15 minutes. We then constructed a peristimulus time histogram (PSTH) for each cell, triggered on the onset of each of the 5 individual images (Fig. 3B). We defined the image with the largest PSTH as the “primary image.” Typically, the PSTH for the primary image had a positive deflection above the average activity, while the PSTH for all other images had a negative deflection. This occurred because the primary image would elicit a much larger response than all the other images, so that it accounted for most of the average activity. Therefore, when we conditioned on the presence of non-primary images, the activity was below the mean unconditioned activity. On rare occasions, a cell would respond similarly well to two or more images. We excluded those cells from our analysis.

**Figure 3:**
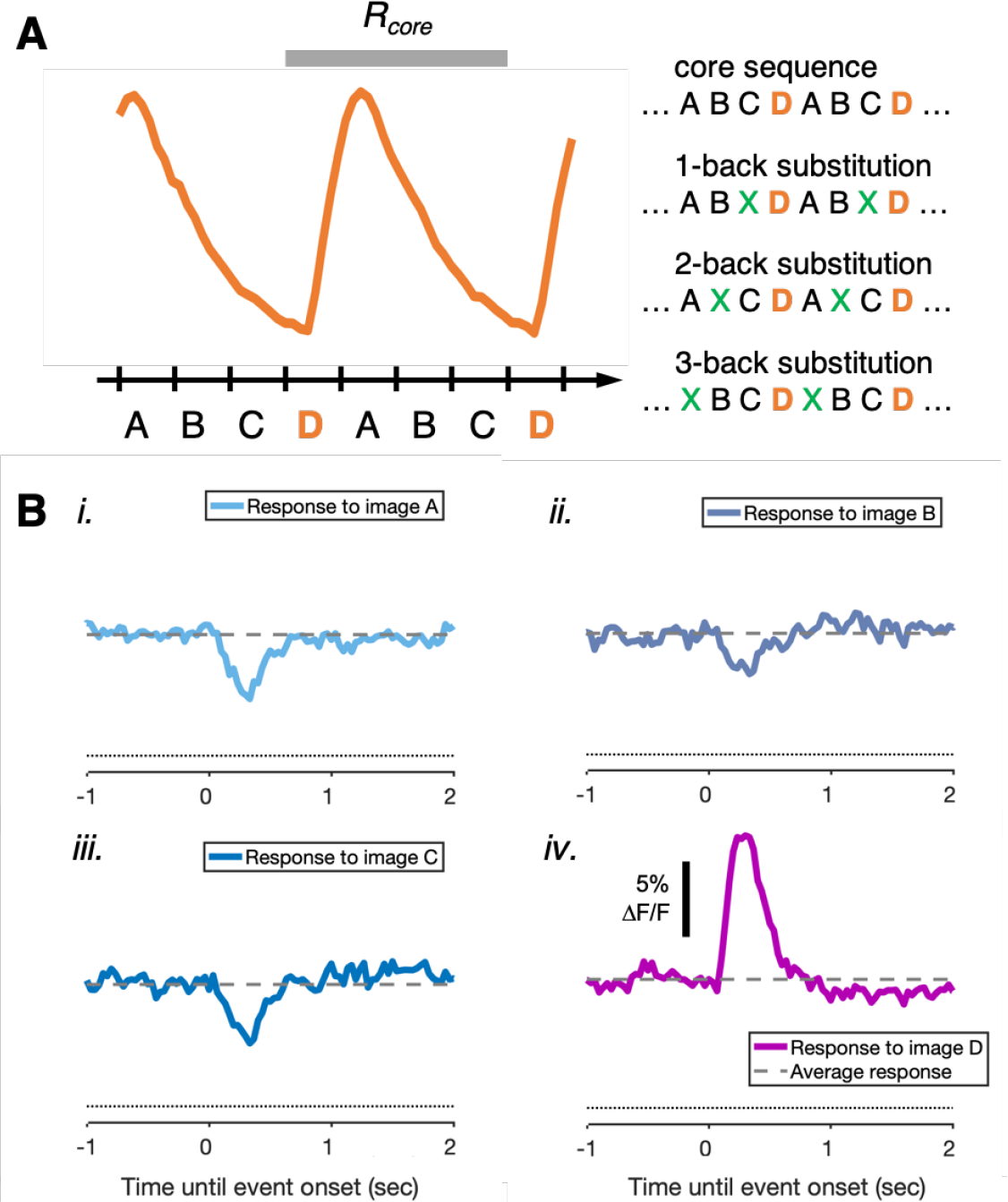
Response characterization of an example cell. (A) Stimulus locked response of an example cell to the core sequence *ABCD* averaged over many trials. Time window used to measure the response amplitude to the core sequence, *R*_core_, shown in grey. (B) Individual PSTHs to images *A*, *B*, *C*, and *D* of the example cell in (A). These PSTHs were calculated during a 15-minute presentation of the 5 images *A*, *B*, *C*, *D*, and *X* in random order. The cell shows an elevated response to image *D*. Images *A*, *B*, and *C* caused a reduction of activity below the mean activity during the entire stimulus ensemble (dashed line). Dotted line indicates baseline activity (∆*F/F* = 0).

In order to confirm our identification of the primary image, we also examined the PSTH during the repeated core sequence. Visual stimuli trigger a response in visual cortical neurons with a delay of about 50-80 msec [33]. The calcium indicator causes an additional delay of 50-80 msec [25]. That means that the response of a neuron to an image would begin to exceed baseline at roughly 100-160 msec after stimulus onset. Taking this delay into account, we were readily able to determine which image caused the largest response (see Fig. 3A). An additional check was to observe a substantial reduction in response when the primary image was substituted by image *X*. All cells included in our analysis exhibited this feature.

#### 2.4.3 Amplitude calculation and definition of relative response

We estimated the average activity of a neuron by averaging the ∆*F/F* signal over one whole cycle of a sequence (Fig. 3A). Given that we selected for neurons that responded primarily to only one image, this one-cycle activity average closely approximated the response to the primary image alone. This simple measure of response amplitude allowed us to ignore cell-specific response delays and different temporal shapes of the PSTH and also provided the best statistical power. We repeated our analysis with smaller time windows, finding qualitatively similar results but somewhat smaller signal-to-noise ratio.

To compute the relative response triggered by an image substitution, we divided the average activity to the substituted sequence by the average activity to the un-substituted (core) sequence:

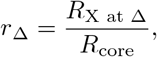

where *R*_X at ∆_ is the average activity to the substituted sequence and *R*_core_ is the average activity to the unsubstituted sequence (Fig. 3A).

To determine if the relative response was significantly different than unity (the null hypothesis), we used the Wilcoxon rank sum test. Then to calculate the error bars displayed in our graphs, we found the standard deviation of a Gaussian distribution having the same p-value.

To characterize the strength of the substitution effect at the population level, we calculated the typical response ratio, 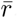, which was given by:

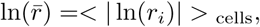

where *r*_*i*_ is the response ratio of cell *i*. This quantity measures the geometric mean of how the response ratio differs from unity, either up or down.

## 3 Results

### 3.1 Layer 2/3 cells are sequence selective

In order to test if neural responses depend on stimulus history, we constructed a sequence of 4 images *ABCD* which we presented to awake, passively viewing mice (Fig. 1 and Methods). These images consisted of random superpositions of Gabor patches, similar in size to V1 receptive fields, and were shown for 250 msec each (Fig. 2). We ran many trials by repeating this “core” sequence. This allowed us to establish a precisely stimulus-locked, averaged response to the core sequence. For further analysis, we then selected cells that showed a strong response to only one image of this core sequence, but not to others (see Methods).

We determined the identity of the image causing the response by constructing PSTHs to the four individual images *A*, *B*, *C*, and *D* from a separate recording with images presented in random order (Fig. 3B and Methods). After establishing the average response to the core sequence, we systematically substituted the images prior to this primary image with another image *X* randomly drawn from the same distribution. This generated sequences *XBCD*, *AXCD*, *ABXD*, and *ABCX*. Cells with a response to image *X* were further excluded from our analysis (see Methods). Comparing the responses to all four substitutions, we found that the timing and basic shape of the response to the primary image remained similar. But interestingly, we found that the amplitude of the response to the primary image was modulated in some cells.

Figure 4A shows this effect in an example cell that responded primarily to image *A*. Panels *i* and *ii* show the response to the core sequence. While responses to individual trials showed variability, mean responses could be constructed with low error bands (Fig. 4A*iii*). When image *A* was substituted with image *X*, the response of this neuron was reduced nearly to zero. This fact is consistent with the hypothesis that the neuron’s response was driven by the immediate presence of image *A*, without significant history effects. In addition, substitution of the previous image *D* (1-back) resulted in no change (Fig. 4A*iv*), again consistent with a lack of any history effect.

**Figure 4:**
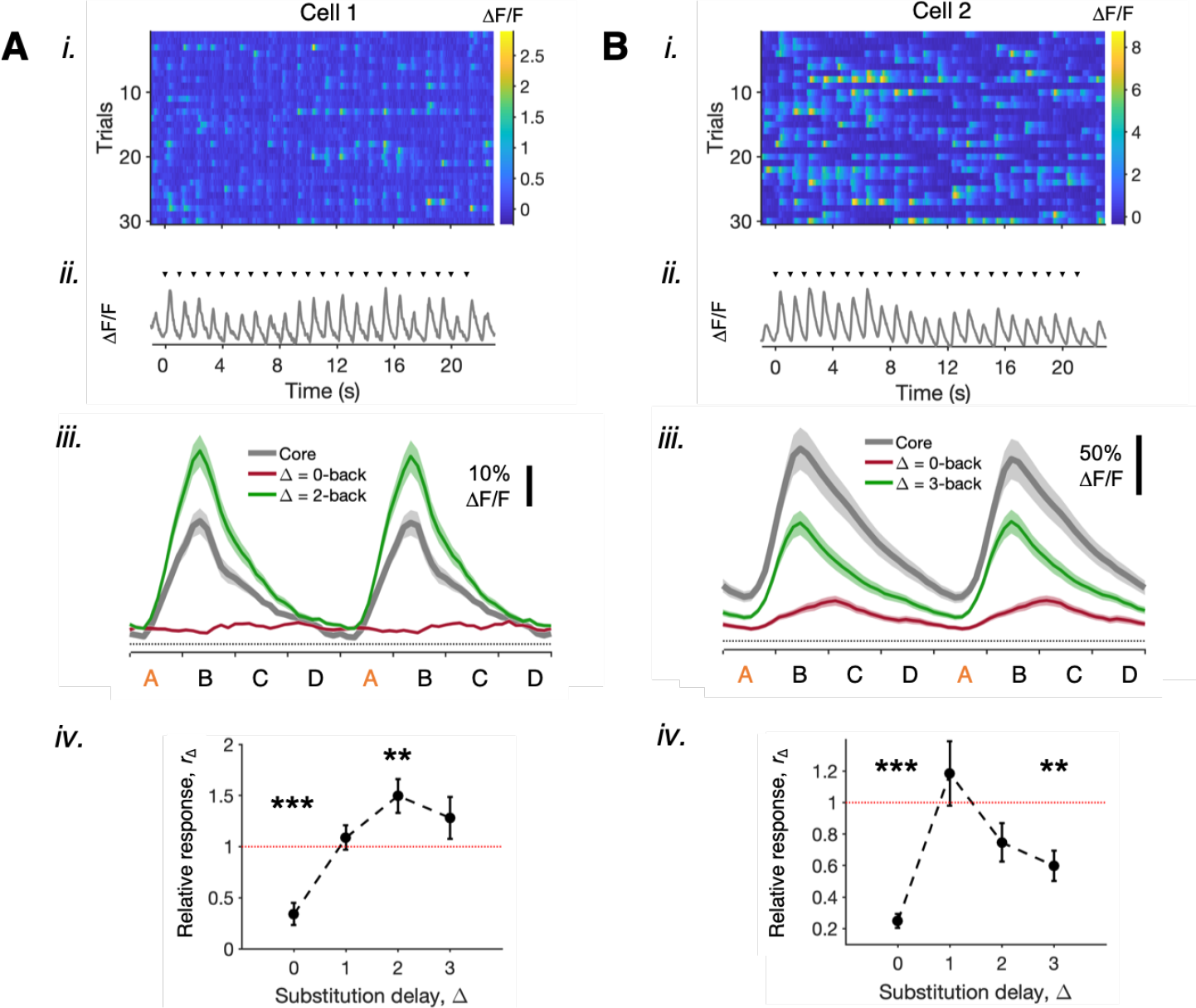
Modulation of the primary image response to different sequence substitutions in the “block” stimulus paradigm. (A) and (B) show two example cells. *i*. Trial-by-trial responses to 30 individual blocks of the repeated core sequence. Calcium activity is represented in color. *ii* Trial-averaged response. Ticks indicate the start of each sequence. *iii*. PSTHs to two cycles of the core sequence *ABCD* (grey), the sequence where the primary image was substituted *XBCD* (red, 0-back), and a sequence where another image was substituted (green, 2-back or 3-back). *iv*. Amplitude of the responses to all possible substitutions (*XBCD*, *AXCD*, *ABXD* and *ABCX*). Statistical significance of each of the four data points, determined by the Wilcoxon rank sum test, with *** indicating p < 0.001 and ** indicating p < 0.01.

However, when image *C* was substituted with *X*, we found a significantly larger response (significance assessed by the Wilcoxon rank sum test). This result demonstrates that, contrary to the previous observations, this neuron was indeed sensitive to the longer time-scale history of the stimulus, in this case the identity of an image presented at a delay of 2 images or 500 msec in the past. All of these observations are consistent with the interpretation that this history effect is *modulatory*. In other words, the presence of image *D* 500 msec in the past did not drive a super-threshold response in this neuron regardless of the currently displayed image. Instead, it modified the response to the primary image, which was capable of driving the neuron above the firing threshold by itself.

Figure 4B shows another example neuron whose primary image was also *A*. When image *A* was substituted with image *X*, the response of this neuron was reduced substantially. This observation is again consistent with our identification of *A* as the primary image. However, when image *B* in the core sequence was substituted with the same image *X* (3-back), we found a significantly smaller response. This change in response revealed an effect of the stimulus history 3 images, or 750 msec, in the past. We quantified the significance of substitutions at all possible image positions for each neuron (Fig. 4A,B *iv*). For both neurons, substitution of the primary image reduced the response, and this reduction was significant. For other substitutions, both neurons exhibited a significant history effect at only one substitution position.

### 3.2 Statistics of the history effect

In order to survey the history effect in the entire population, we plotted a rank-ordered list of the relative response for all cells, separated by the substitution delay (Fig. 5A). We found that substitutions increased the primary image response of some cells by a factor as large as 3; other substitutions decreased the response by factors up to 2. Furthermore, from a statistical point of view cells were less affected by substitutions that were further back in time, consistent with the basic intuition that the history effect should die out for sufficiently long delays. This can be seen as a subtle shift of the distribution of relative response values closer to unity for longer delays. Indeed, we calculated the typical response ratio, *r̄*, which measures the factor by which the relative response differs from unity (see Methods). Its value was 1.8 for 1-back substitutions, and it decreased steadily for increasing substitution delays (Fig. 5D).

**Figure 5:**
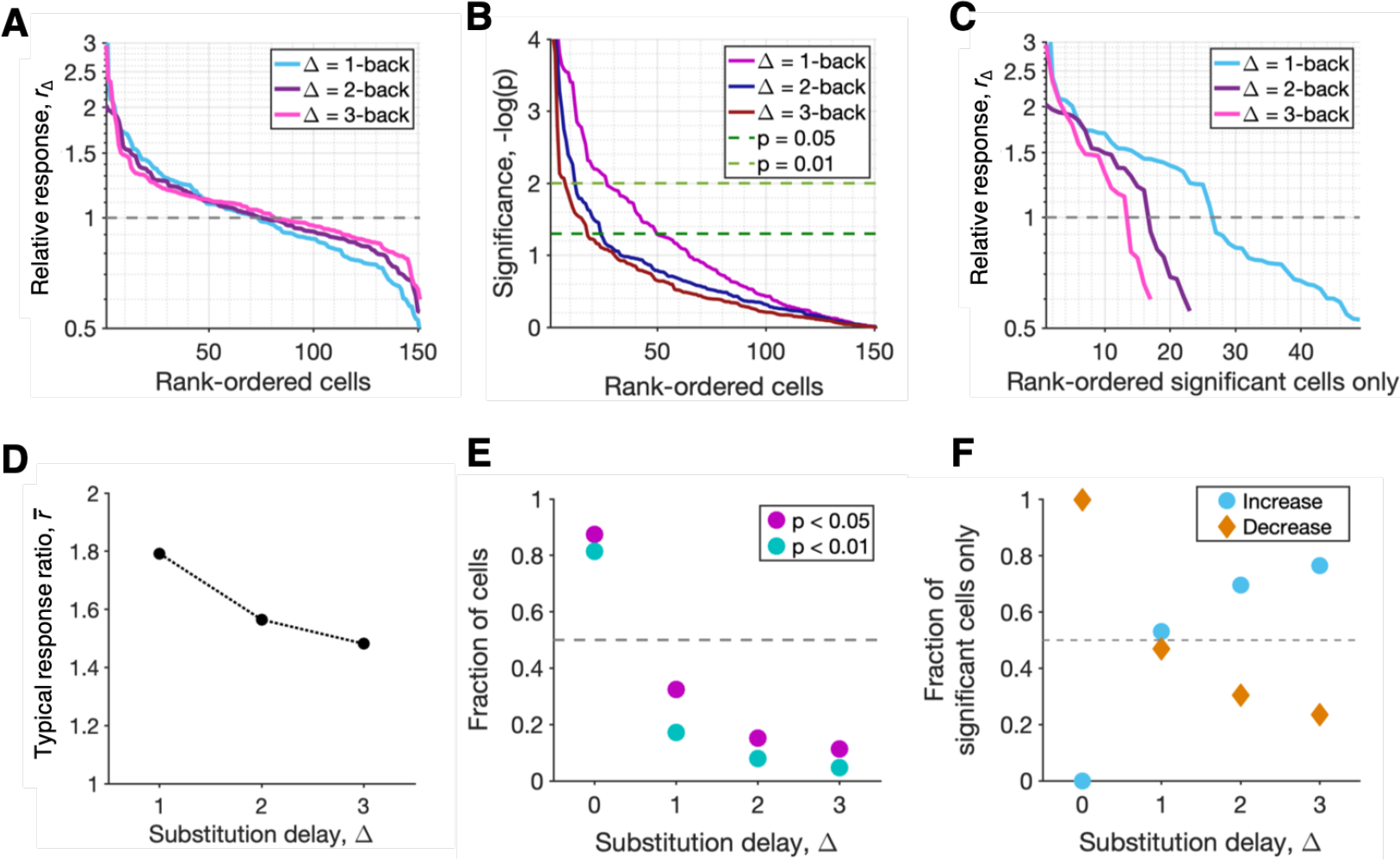
Population-level statistics of the history effect. (A) All 151 cells ranked by relative response amplitudes for each substitution delay (different colored lines). (B) All cells ranked by the significance of the modulation of the primary image response for each substitution delay (different colors). (C) Rank-ordered relative response of all neurons showing a significant history effect. (D) Typical response ratio as a function of the substitution delay. (E) Fraction of cells showing a significant history effect plotted versus substitution delay. (F) Fraction of cells with an increased response due to the substitution (relative response *>* 1) or decreased response (relative response < 1) at each substitution delay.

Many of the relative responses shown in Fig. 5A were not statistically significant, so we also plotted a rank-ordered list of the statistical significance of each relative response (Fig. 5B). Here, we found a larger fraction of significant effects for shorter substitution delays, again consistent with a systematically weakening effect at longer delays.

Restricting our analysis to statistically significant modulations in response only, we found that the majority of cells exhibited larger responses due to the substitution image (Fig. 5C; relative response *>* 1). It should be noted, however, that significance is more difficult to establish for a decrease in activity because the spontaneous activity of a particular cell generates an underlying “floor” of noisy activity (see Fig. 4A*iii* and B*iii* for two example cells exhibiting a certain level of spontaneous activity). Plotting the fraction of significant history effects versus delay (Fig. 5E), we found that roughly a third of all cells showed a significant change in activity to a 1-back substitution (29%, p < 0.05), but only 12% of all cells showed a significant change in activity to a 3-back substitution. Among all the significantly modulated cells, we found that substitutions further in the past were more likely to produce a larger response to the primary image compared to a smaller response (Fig. 5F). Finally, we note that while the history effect weakened with longer delays, it clearly had not decayed away entirely. This motivated us to carry out experiments with longer sequences to see how far back into the past the effect extended.

### 3.3 History effects extend up to 5 images in the past

In order to see how far back the history effect extended, we performed experiments with longer sequences. Figure 6A shows an example cell whose primary image was *C* in the 8-image sequence, *ABCDEFGH*. When the primary image was replaced with image *X*, the response was again reduced to nearly zero. What was remarkable, however, was that when image *F*, which was five images back from *C*, was replaced there was a significant reduction in response. None of the other substitution positions produced a significant response reduction (Fig. 6A*ii*). Figure 6B shows an example cell whose primary image was *O* in the 16-image sequence, *ABCDEFGHIJKLMNOP*. Substitution of the primary image again reduced the response nearly to zero. When image *K* – four back from the primary image – was substituted, however, we observed a significant increase in response. While most substitutions produced relative responses greater than 1, only the substitution at 4 images in the past was significant (Fig. 6B*ii*). These examples show that the modulatory effect of stimulus history tended to be highly selective for a particular substitution delay.

**Figure 6:**
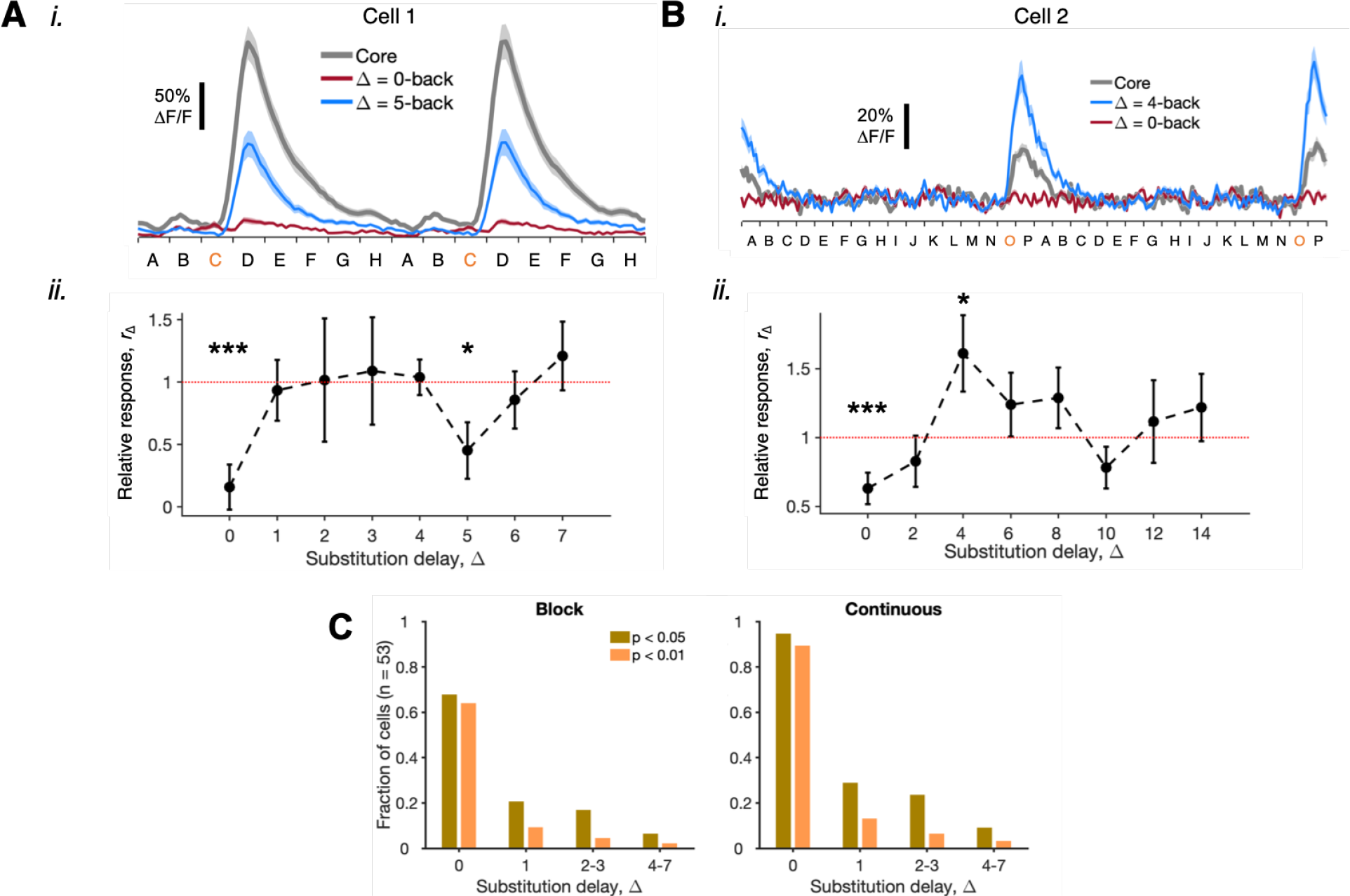
Experiments with 8- and 16-image sequences. (A) *i*. PSTHs of an example cell responding to an 8-image sequence: core sequence *ABCDEFGH* (grey), the primary image substitution *ABXDEFGH* (red) and a substitution 5 images back from the primary image *ABCDEXGH* (blue). *ii*. Amplitude of the responses to all possible substitutions (*XBCD*, *AXCD*, *ABXD* and *ABCX*). Statistical significance of each relative response value was determined by the Wilcoxon rank sum test, with *** indicating p < 0.001 and * indicating p < 0.05. (B) A different cell responding to 16-image sequences. (C) Fraction of cells showing significant modulation of the response to their primary image. Brown: p < 0.05, orange: p < 0.01. *Left*: “block” stimulus design, *Right*: “continuous” stimulus design. For both designs, we found similar proportion of cells with a significant modulation.

We then quantified this selectivity for substitution delay and found that a majority of the cells were only selective to one particular substitution delay. This result was found for all sequence lengths tested: for 4-image sequences, 44 cells (67.7%) were only selective to a single substitution delay; for 8-image sequences, the fraction was 62.5%; for 16-image sequences, the fraction was 61.9%.

### 3.4 Stimulus paradigm does not affect the extent of sequence selectivity

In the block design of our experiment, we found elevated activity in most neurons for the first few presentations of a new image sequence. This population-level phenomenon has previously been called the *transient response* [26]. To investigate whether the transient response played a role in shaping sequence selectivity, we carried out experiments with a continuous design instead. We then compared history effects with these two stimulus designs by measuring the fraction of all substitutions tested that produced a significant change in the response to the primary image (Fig. 6C). We found consistent results with both stimulus designs (data pooled over *n* = 3 mice). As a result, we pooled the data obtained from these two different stimulus designs for subsequent analysis.

### 3.5 Cell-specific history effects are reproducible across days

In order to study if history effects were stable across days, we repeated the 8-image sequence experiment for two consecutive days. We extracted from our data all cells that showed a significant history effect on at least one day. We then plotted the relative response on day 1 against the relative response on day 2. If the history effect were consistent, then neurons that showed a response increase on day 1 would also show a response increase on day 2, and *vice versa*. To check for statistical significance, we performed a Wilcoxon signed rank sum test to determine whether the fraction of cells with a consistent history effect across both days was significantly greater than chance. We found that history effects were, indeed, consistent across days (Fig. 7). This result suggests that the effect originated from a somewhat “hard-wired” property of the circuit that is preserved across days.

**Figure 7:**
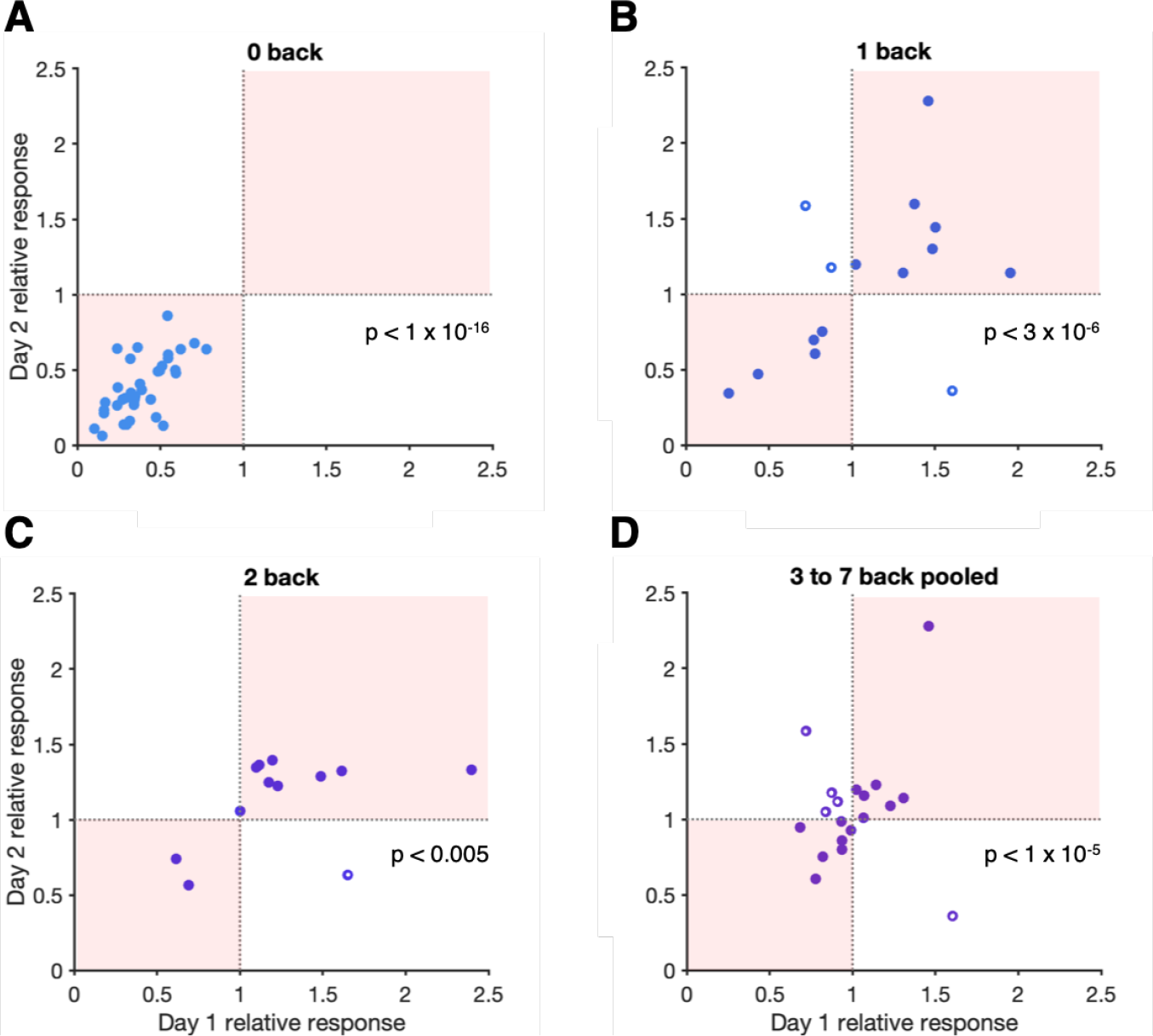
Cell-by-cell reproducibility of modulation effects at different substitution delays taken from the 8-image data. Day 1 vs. Day 2. Consistent history effects are shown by shaded quadrants of the graph as well as by solid circles. Inconsistent effects are shown by open circles. (A) Substituting the primary image. (B) Substituting the image before the primary image (1-back). (C) Substituting the images 2 back. (D) Substituting images 3 back.

### 3.6 Characterizing the history effect in the entire population

To quantify history effects across the entire population of layer 2/3 neurons, we measured the fraction of all substitutions that exhibited a significant effect. To this end, we pooled all data from the block design and the continuous design (see Methods). Data were separated by substitution delay and by the length of the core sequence (Fig. 8). Experiment-level significance was calculated using Poisson statistics. For four-image sequences (*n* = 151 neurons), we found that history effects were highly significant for all delays, at both the p < 0.05 and < 0.01 thresholds (Fig. 8A). For 8- and 16-image sequences, the fraction of substitutions showing a significant effect was consistent with that measured with 4-image sequences (Fig. 8B). We also found a significant effect up to 4-5 images into the past (data combined), showing an impact of an image presented over 1 second before the primary image.

**Figure 8:**
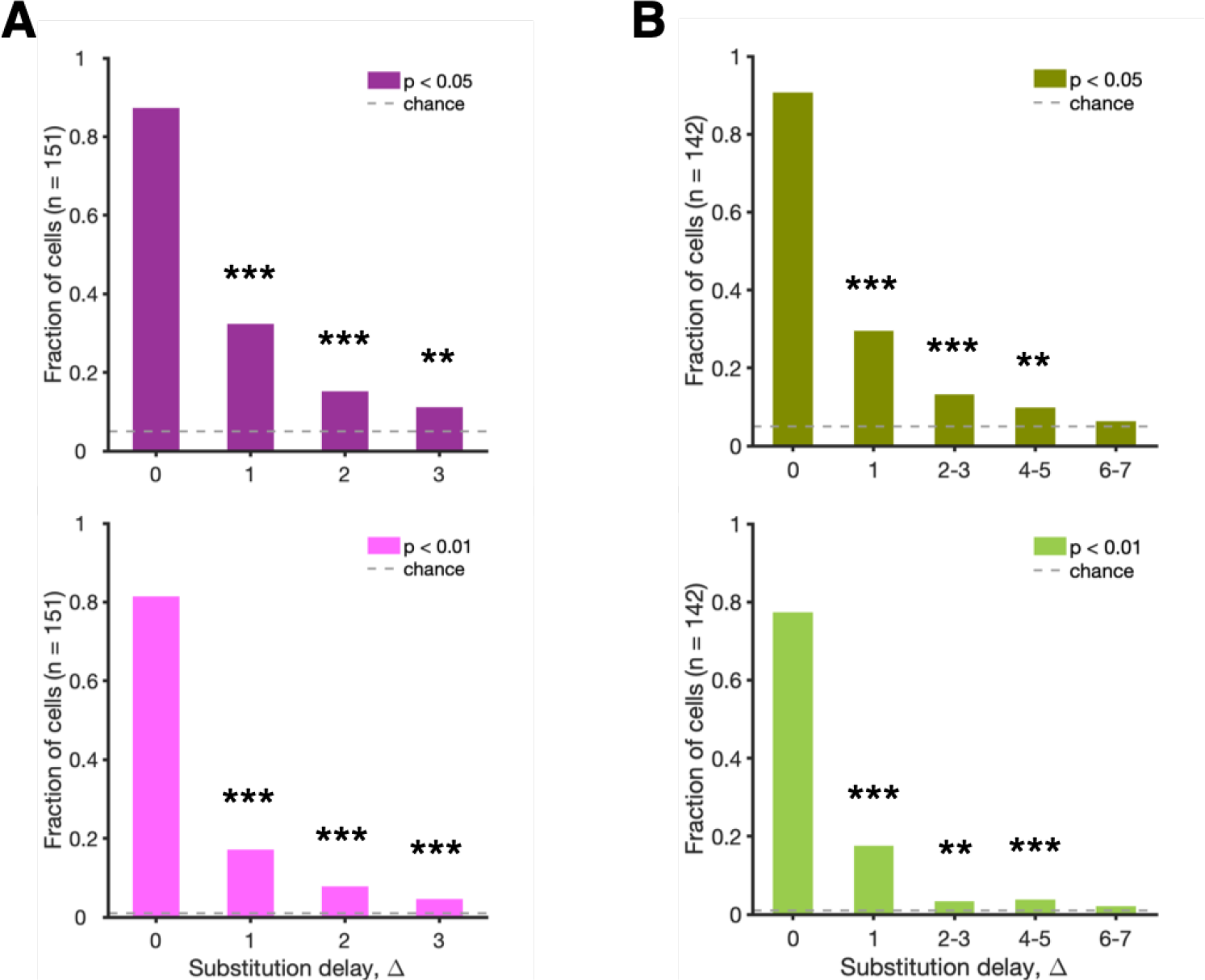
Fraction of cells showing a significant modulation of the response to their primary image as a function of substitution delay, ∆. (A) 4-image sequence data. (B) 8- and 16-image sequence data combined. (*Upper*, *lower*) panels use experiment-level significance thresholds of (p < 0.05, p < 0.01).

## 4 Discussion

In this study, we have demonstrated a new form of short-term memory at the level of single neurons in the primary visual cortex. Our methodology was first to stimulate neurons with a core sequence of images. Under these stimulus conditions, the vast majority of recorded neurons had a strong response to a single, primary image within the core sequence. Then, we substituted images in the core sequence with a new, standard image. We observed significant changes in the response amplitude of V1 neurons to the primary image when previous images were substituted with a different image drawn from the same image ensemble. These effects were heterogeneous across cells and present at a sparse subset of time intervals. However, roughly half of all tested neurons showed a significant history effect for at least one substitution delay (for 4-image experiments, 43% of all cells had a history effect; for 8-image experiments, 62% had a history effect).

Due to our exclusion criteria, all of our analyzed neurons exhibited a reduction in response when we substituted the primary image. However, the relative response was typically in the range of 0.1 – 0.2, rather than zero. This non-zero relative response could be due to several factors. First, neurons exhibited a baseline level of activity that was due to spontaneous firing or activity not locked to our external stimulus events. As we did not attempt to subtract this baseline, this activity would be present both for the core sequence and the substitution.

Second, neurons could have a weak response to the substitution image, *X*. Third, the history of stimuli before the primary image could, in principle, be strong enough to drive spiking responses in the neuron. Regardless of the source of remaining activity when we substituted for the primary image, we found that a large fraction (261 out of 293 cells = 89%) but not all neurons had a significantly reduced response. The fact that some neurons had a reduction that did not reach statistical significance presumably relates to our experimental design (*i.e*., a combination of the noise level with our choice of number of trials).

For substitutions that cause an increase in the response to the primary image, we interpret the original stimulus history effect to be *suppressive*. This is because our substitution eliminates the particular history effect due to the original image. Similarly, substitutions that cause a decrease in response to the primary image suggest that the original history effect was *facilitatory*. This interpretation has some caveats. First, it assumes that the substitution image, *X*, did not generate any response in the recorded neuron by itself. While we screened out cases of strong responses to image *X*, weak responses could remain. This confound would increase the apparent response to the primary image, thereby leading to larger relative responses. Therefore, this effect would tend to raise the number of substitutions that caused a response increase versus those that caused a decrease. This factor might play a role in the greater abundance of response increases that we observed (Fig. 5E).

Second, this interpretation also assumes that the new image *X* does not trigger its own history effect. If the history effect was due to image *X* itself, this would change our interpretation – *i.e*. an increase in response to the primary image would imply a facilitatory history effect and *vice versa* – but would not invalidate the observation of a significant history effect. Furthermore, the relative rarity of history effects suggests that this confound should also be rare. A future direction might be to test with more than one substitution image to see if effects are consistent, or perhaps substitute a blank image, which should drive much less activity by itself.

A change in response to the primary image due to substitution of the previous image (*i.e*., a 1-back history effect) should be viewed as a special case, because this phenomenon has a relatively simple explanation. Sensory neurons exhibit temporal integration, which can be characterized by reverse correlating between neural activity and a white noise stimulus [34]. This analysis typically reveals integration over 100–200 msec for neurons in V1 [35][36][37]. Thus, a 1-back history effect is consistent with the typical time scale of temporal integration. Given this simple mechanism, it is perhaps somewhat surprising to observe such a *small* fraction of neurons with 1-back history effects (~28%; Fig. 8). In any case, history effects at least 2 images in the past require a different mechanism, as the duration of each image (250 msec) implies that temporal integration would be unlikely to provide an explanation. Furthermore, the computational implications of longer history effects are potentially more far-reaching. Such effects were robustly present in a large fraction of all neurons tested (for 8-image experiments, 51 out of 91 = 56% of all cells had a history effect at 2 or more images into the past).

Several different circuit mechanisms could contribute to our findings. First, a past image, *A*, could activate decaying activity that outlasts image *A* in a subset of neurons that are selective for that image (*i.e*. differentially activated by the substitution image, *X*). Of course, the layer 2/3 neurons that we recorded typically did not show this form of activity, instead firing strongly only to the primary image. But such decaying, selective activity could be present in unrecorded neurons that serve as the input to our recorded neurons. We will call these un-recorded, presynaptic neurons, ‘subunits’. Such subunits could then directly excite the recorded neuron, producing a facilitatory history effect, or activate feedforward inhibition, producing a suppressive effect (Fig. 9A). When we substitute image *X* for image *A*, a different subpopulation of unrecorded neurons would be activated (not shown in Fig. 9), and so the decaying activity in subunits produced by image *A* would be changed. As a result, the response of the recorded neuron to the primary image would change. The decaying activity of these subunits would give rise to a history effect that would exist over a broad range of time scales. However, each image triggers a different subset of subunits with decaying activity (*e.g*., red versus blue cells in Fig. 9A), so that our substitution experiment would still exhibit temporal specificity.

**Figure 9:**
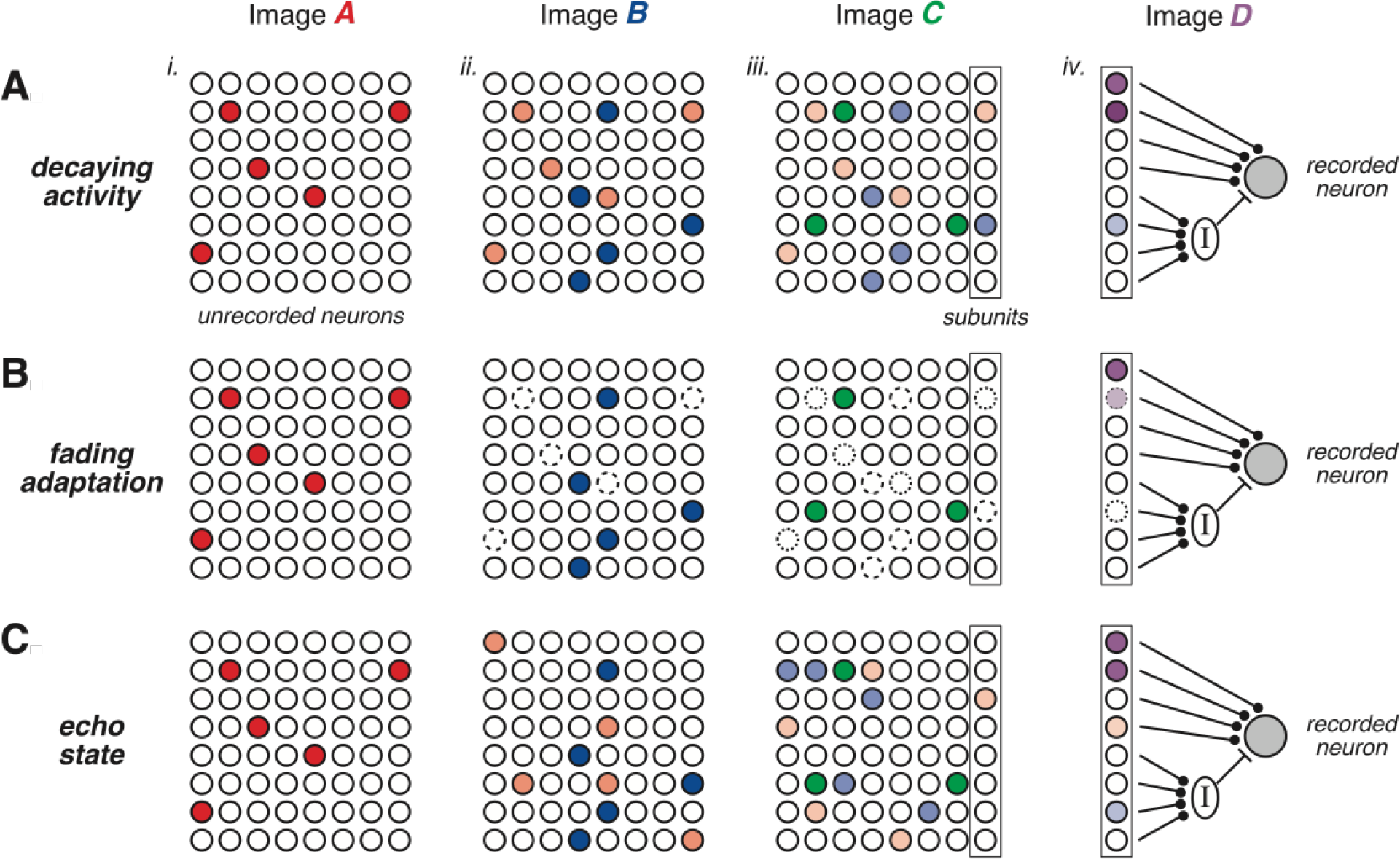
Circuit-Level Models of the History Effect. (A) *i*. In the decaying activity model, an unrecorded neural population (small circles) is sparsely activated by image *A* (red). *ii*. Previous activation decays slowly (lighter red) while image *B* activates a new sparse population (blue). *iii*. Previous activity decays (lighter colors) while image *C* activates yet another sparse population (green). Neurons in the right column (boxed) are presynaptic to the recorded neuron (subunits). *iv*. Subunit activity combines decaying previous activity along with new excitation from image *D*. Subunits activate recorded neuron (grey) either through direct excitation or indirect inhibition via an inhibitory neuron (I). Direct pathways produce a facilitatory history effect, and indirect pathways produce a suppressive effect. (B) Similar diagram for the fading adaptation model. Here, the memory of previous activation is not expressed by decaying activity (circles are open) but by an internal adaptation variable (dashed line of circle). Now when the primary image, *D*, triggers subunits, their response depends on adaptive state (light purple). (C) Similar diagram for the echo state model. Here, decaying activity is expressed in new subpopulations in each time step.

A second model involves fading effects of adaptation in unrecorded neurons. Here, the past image could activate a neuron, triggering short-term adaptation, like depression, which would last beyond the immediate time scale of the neuron’s response. Then, when the primary image was displayed, this subunit would provide a decreased input to the recorded neuron, compared to its response if a different history had occurred (Fig. 9B). This would produce a suppressive history effect. Alternatively, this subunit could experience facilitation due to the earlier image or activate the recorded neuron via feed-forward inhibition. In both of these cases, the subunit would provide increased input to the recorded neuron, thus creating a facilitatory history effect. Notice that such a subunit would need to be co-tuned to the original image, *A*, and the primary image, *D*. Similar to the decaying activity model, this *fading adaptation* model implies history effects that would be broadly distributed across time, yet would still produce temporally specific perturbations in our substitution experiments.

A third model involves an echo state network or liquid state machine [23][38]. In this model, the earlier image triggers one subpopulation of neurons. Then, the intrinsic dynamics of the local recurrent neural network would lead to the activation of different subpopulations at successive steps forward in time. Due to noise and/or deterministic chaos, these changing subpopulations, or echoes, would fade over time (Fig. 9C). Because we typically did not see significant activation when we substituted for the primary image, these echoes would need to be present in subunits rather than recorded neurons. As before, these subunits could directly excite the recorded neuron, producing a facilitatory history effect, or provide disynaptic inhibition, producing a suppressive effect. If instead the earlier image *A* was substituted for *X*, a different sub-population of echoing neurons would be activated, and as long as this subpopulation provided different inputs to the recorded neuron, there would be a change in its response to the primary image.

The *decaying activity* and *fading adaptation* models are both similar. The main difference between them is that in the former, the memory of a previous stimulus is encoded by active firing of neurons, while in the latter, the memory is encoded in the adaptive state of neurons, which is not directly observable in our recordings. This makes the *fading adaptation* model seem more likely, both because we don’t see any evidence of decaying activity that outlasts the presentation of a neuron’s primary image and because adaptation has been widely reported in V1 while decaying activity has not been. There is a related challenge to the *echo state* model, namely that firing of subpopulations of neurons is required to sustain echoes across time. Such echoes would be expected to drive activity of recorded neurons, even when we substituted for their primary image. The fact that we could only confidently observe modulatory history effects, makes the *echo state* model less likely. However, we cannot rule out the possibility that the echoes are comprised of dense subpopulations of weakly firing neurons rather than a sparse subpopulation of strongly firing neurons, as depicted in Figure 9.

The *echo state* model differs from the other two in that the history effect induced by any given image is temporally specific, while in the other two models the history effect should be broadly similar across time. This implies an experiment that can potentially distinguish the *echo state* model from the other two. In this experiment, we could present an earlier image, like *A*, at different time intervals before the primary image (*D*). In the echo state model, the history effect would be completely different at different time intervals, while in the other two models, the history effect should have the same polarity – either facilitatory or suppressive – albeit with different strength.

Several kinds of history effects have been reported in the literature. Some, like prediction suppression [2] require a multi-day learning process to create. These phenomena embody a form of memory that is *selective*, in the sense that only trained associations will be encoded in the system. The history effects that we report here, in contrast, are immediately present for all stimuli tested. In this sense, our results imply a form of memory in the system that is *general*. Selective and general memory systems each have their own benefits and drawbacks, so it is not surprising that both are present in neural circuits.

Most previous studies of general short-term memory phenomena have reported relatively short time scales, typically 100-200 msec. Our study demonstrates a significantly longer memory time scale, up to ~1000 msec. This increase in time scale is significant, because there are more profound computational implications of a longer time-scale general memory than a shorter one. However, it should be noted that both the auditory and somatosensory pathways have shorter response latencies and somewhat faster intrinsic time scales. Thus, if temporal dynamics in different sensory systems are scaled with respect to their “core processing time”, our results in the visual system might not be so different from reports in those of other sensory systems.

The study by Nikolić *et al*. [22] did find history effects with a long time-scale, comparable to our results. One difference is that this study used a population decoding strategy to demonstrate a history effect, rather than analysis of single neurons. While these two approaches are complementary, our analysis revealed properties that begin to give some insight into the circuit mechanisms. In particular, our results challenge the echo state model, because our history effects appear to be modulatory. Another difference between the studies is that the fading activity shown in Nikolić *et al*. was substantially disrupted by the presentation of a second image. In contrast, we presented sequences of images with no blank periods in between, so that all of our history effects occurred in the context of interference from new visual responses. This robustness to interference also seems most consistent with the *fading adaptation* model.

## Acknowledgments

The authors thank the members of the Berry Lab for helpful comments on the manuscript, members of the Tank Lab for technical assistance, and especially Sue Ann Koay for guidance on describing the procedure of widefield imaging. The work was supported by a grant EY02675 from the National Eye Institute and a grant from the PNI Innovation Fund.

